# Impaired synaptic inhibition and enhanced aversion encoding by lateral habenula neurons during Δ^9^-tetrahydrocannabinol withdrawal

**DOI:** 10.1101/2025.11.24.690244

**Authors:** Eun-Kyung Hwang, Deborah Daphne, Charlie C. Maddox, Agustin Zapata, Alexander F. Hoffman, Carl R. Lupica

## Abstract

The lateral habenula (LHb) integrates cortical and basal forebrain (BF) input to control monoaminergic outflow and is implicated in depression, anxiety, impulsivity, and aversion. Although cannabis withdrawal is associated with negative affect and heightened anxiety, LHb involvement is not established. Here, effects of withdrawal from the psychoactive cannabis constituent Δ^9^-tetrahydrocannabinol (Δ^9^-THC) on LHb neurons were assessed with photometric calcium measurement during fear conditioning and *in vitro* electrophysiology. LHb calcium signals were larger during footshock, and presentation of a tone paired with footshock in Δ^9^-THC withdrawn rats. Fear-induced freezing to the tone was also larger during Δ^9^-THC withdrawal. Electrophysiology revealed larger LHb excitatory-inhibitory (E-I) ratios during Δ^9^-THC withdrawal, resulting from impaired synaptic GABA release. Moreover, GABA release via optogenetic activation of BF-LHb inputs was impaired during Δ^9^-THC withdrawal, whereas no changes occurred at ventral tegmental area-LHb inputs. Recovery of BF-LHb GABA release and cannabinoid CB1 receptor desensitization were incomplete at 30 days of Δ^9^-THC withdrawal. The data show that LHb responsivity to primary aversive and conditioned stimuli are increased during Δ^9^-THC withdrawal, and that this is likely mediated by altered E-I balance. We propose negative affect experienced during cannabis withdrawal is related to LHb hypersensitivity to aversive stimuli and this facilitates encoding of associated environmental cues.

## Introduction

Addictive drug use is thought to be initially driven by the desire to experience pleasurable effects that include euphoria, elevated mood, and a general state of well-being, via interactions with brain reward systems [1]. However, long-term drug use promotes adaptation of neural circuits leading to the development of dependence that is manifest as a withdrawal state when drug use ceases. Withdrawal from several drugs, including opioids, alcohol, psychostimulants, nicotine, and cannabis, is associated with both physiological and psychological symptoms that include increased sensitivity to stress, irritability, heightened anxiety, and a general negative dysphoric state [2]. Moreover, avoidance of these negative withdrawal symptoms through continued use of the drug is considered a primary obstacle to maintaining drug abstinence [1–4]. Efforts to identify brain regions involved in drug dependence and negative withdrawal states have implicated the ventral striatum, extended amygdala, periaqueductal gray, cingulate cortex and median raphe nucleus [2,5,6]. However, another brain structure that is involved in processing aversion and negative emotional states, known as the lateral habenula (LHb), has more recently been implicated [7–11].

Consistent with its role in processing negative affect, LHb neuronal activity increases in response to situations and stimuli associated with negative outcomes, such as the absence of expected rewards (reward prediction error, RPE) [12], or the presence of aversive stimuli. In contrast, LHb activity decreases when unexpected rewards are received [13]. Importantly, LHb neuron activity is 180 degrees out of phase with the activity of the midbrain dopamine (DA) neurons that encode reward [10,12].

LHb output neurons are largely glutamatergic [14–16], and evidence suggests that LHb involvement in negative affective states is mediated through its control of monoaminergic neurons, such as DA cells in the ventral midbrain and hindbrain serotonin (5-HT) cells that extensively innervate forebrain areas [7,10,17,18]. Although direct connections between the LHb and these monoaminergic regions exist [14,19], the inhibition of DA and 5-HT neurons by the LHb during negative affective states occurs via excitation of GABAergic rostromedial tegmentum (RMTg) neurons that target the monoaminergic cells [19–25]. These studies, together with those showing that LHb neurons are sensitive to opioids, cocaine, alcohol, nicotine and cannabinoids, has led to the hypothesis that changes in LHb activity may underlie some acute aversive effects of these drugs, as well as the negative affective states observed during withdrawal from long-term exposure [2,6,11,17,26–30].

Although strong evidence supports the idea that drug withdrawal is associated with negative physiological and psychological states, only recently has cannabis use disorder (CUD) and the existence of a cannabis withdrawal state been codified in humans in the Diagnostic and Statistical Manual of Mental Disorders (DSM-5) [31,32]. Thus, cessation of long-term exposure to Δ^9^-THC and other cannabinoids results in behavioral withdrawal symptoms and changes in brain function similar to that observed for other addictive drugs. For example, diminished forebrain DA function during drug withdrawal is observed with opioids and cocaine, as well as cannabis [28,33–35]. More generally however, the consequences of cannabis withdrawal on brain function is poorly understood compared to that of other psychoactive drugs.

Recent studies demonstrate functional cannabinoid CB1Rs in the LHb, where they regulate synaptic transmission and behavior [26,36–38]. Moreover, withdrawal from long-term Δ^9^-THC exposure has recently been shown to reduce LHb neuron spontaneous firing, supporting the idea that the LHb and its connections with other brain regions may be involved in behavioral changes seen during cannabis withdrawal [28]. Despite these advances in our knowledge of the consequences of CUD and the neurobiology of cannabis withdrawal, the mechanisms through which LHb function may be altered and how this may modify behavior remain poorly understood. Here, we explore the hypothesis that synaptic inputs controlling LHb neuron excitability are altered, and that this contributes to heightened neuronal activity and behavioral responses to aversive stimuli during Δ^9^-THC withdrawal.

## Results

### Increased footshock-evoked LHb neuron calcium signal during Δ^9^-THC withdrawal

To determine whether the sensitivity of LHb neurons to aversive stimuli is altered during Δ^9^-THC withdrawal we used in vivo fiber photometry to measure GCaMP8f-mediated calcium responses during Pavlovian fear conditioning (FC). A mild footshock unconditioned stimulus (US, 0.5 sec, 0.6mA) was paired with an auditory tone (CS) while GCaMP8f calcium responses were measured, beginning 2-d after drug or vehicle withdrawal (Veh, **Fig. 1A**). FC (CS-US association) was conducted in 10, 20-sec trials (**Fig. 1B-C**) on 2 consecutive days (FC1, FC2; **Fig. 1A)**. Consistent with previous reports in naïve rodents [39,40], footshock was associated with a significant increase in LHb calcium signal in both treatment groups during FC1 (**Figs. 1B-D**; 2-way ANOVA, F_9,170_ = 8.67, p<0.0001, main effect of trial). However, the calcium signal increase was significantly larger in the Δ^9^-THC withdrawal group (**Fig. 1B-D**, 2-way ANOVA, time x treatment interaction, F_9,170_ = 3.38, p=0.0009, and treatment main effect = F_1,170_ = 62.762, p < 0.0001), with differences seen on trials 1-3,6-7 (p=0.0041, p<0.0001, p=0.021, p<0.0001, p=0.017, respectively, Šídák’s post-hoc). The groups did not differ upon US presentation during FC2 (**Fig. 1E**, top; treatment x trial interaction, F_9,140_ = 0.536, p = 0.847). The data suggests that the excitation of LHb neurons to US presentation was greater in the Δ^9^-THC withdrawal group during the first day of fear conditioning compared to Veh controls.

**Figure 1:**
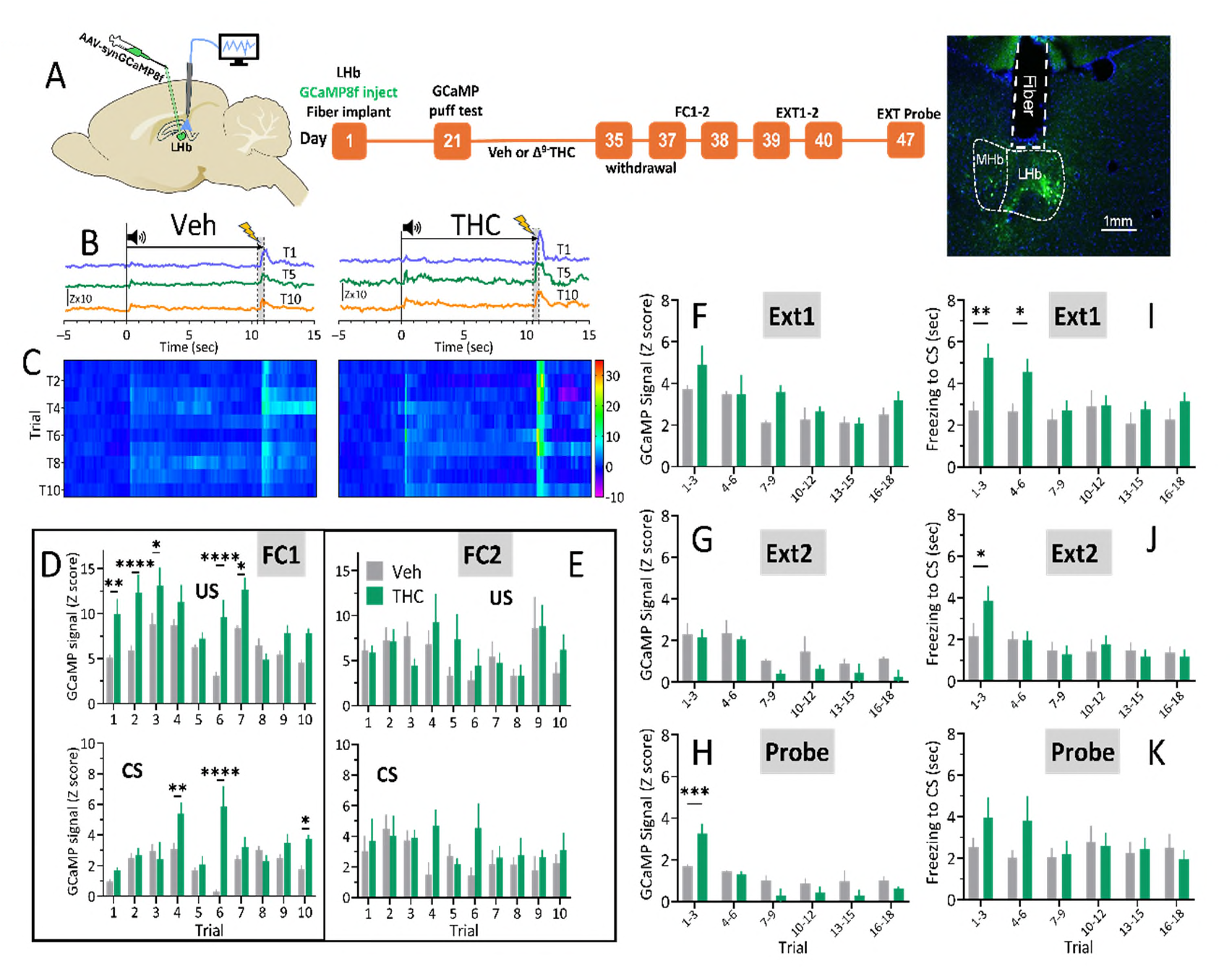
LHb GCaMP8f fluorescence during fear conditioning (FC) and Δ^9^-THC or Veh withdrawal. **A**. Left: Diagram of virus injection into LHb and *in vivo* fiber photometry setup. Center: Experimental timeline for photometry and behavior. Right: Example histology of fiber placement in LHb and GCaMP8f expression. **B**. Mean Z-score changes in GCaMP8f signals during trials 1, 5, and 10 of FC1, 2-d post Veh (left, n= 10) or Δ^9^-THC (right, n = 10) withdrawal. Tone onset at time 0 sec, footshock application at 10.5-11 sec (shaded bar). **C**. FC1: Heatmaps of mean GCaMP8f Z-score changes across all trials during Veh (left, n=12) or Δ^9^-THC (right, n=7) withdrawal (same timecourse as **B**). **D**. Top: GCaMP8f Z-score change during footshock (US) across 10 trials of FC1 at 2-d withdrawal from Δ^9^-THC or Veh. Significant trial x treatment interaction (F_9,170_=3.38, p=0.0009); differences at trials 1-3,6-7 (Sidak’s posthoc); significant main effects of trial (F_9,170_ = 8.67, p<0.0001) and treatment (F_1,170_ = 62.76, p<0.0001). Bottom: Peak GCaMP8f Z-score during tone (CS), 0-1 sec window, during FC1. Significant trial x treatment interaction (F_9,170_ = 7.64, p<0.0001); differences at trials 4, 6, 10 (Sidak’s, ****= p<.0001, **= p<.01, *=p<.05). **E**. FC2: Top, peak GCaMP8f z-score to US. No significant interaction (F_9,140_= 0.536, p = 0.847) or main effects. Bottom: CS responses, no significant interaction (F_9,140_= 0.854, p = 0.568); significant treatment effect (F_9,140_ = 4.19, p = 0.043). **F**. Extinction 1 (Ext1): Peak GCaMP8f Z-score during CS. No trial x treatment interaction (F_5,102_ = 0.995, p = 0.425); significant trial (F_5,102_=6.29, p<0.0001) and treatment main effects (F_1,102_=5.84, p=0.017). **G**. Ext2: GCaMP8f Z-score. No significant trial x treatment interaction (F_5,102_ =0.20, p=0.096), significant trial (F_5,102_=5.11, p=0.0003) and treatment (F_1,102_=3.95, p=0.0496) main effects. **H**. Probe test (1-week post-Ext2): Significant trial x treatment interaction (F_5,90_ =4.80, p=0.0006); Significant GCaMP8f increase on trials 1-3 (p=0.0007, Sidak’s). **I**. Ext1: Time spent immobile (freezing) during presentation of CS. No significant trial x treatment interaction (F_5,132_=1.52, p=0.188), significant trial (F_5,132_=2.82, p=0.019) and treatment (F_5,132_=11.92, p=0.0007) main effects. Differences at trials 1-3, and 4-6, p=0.0012 and 0.0144, Sidak’s (n=11, 13, THC, Veh). **J**. Ext2: Freezing to CS. No significant trial x treatment interaction (F_5,132_=1.35, p=0.246), significant trial main effect (F_5,132_=4.04, p=0.0019), no significant treatment main effect (F_5,132_=0.667, p=0.416). Difference at trials 1-3, p=0.010, Sidak’s. **K**. Probe extinction test: Freezing to CS. No significant trial x treatment interaction (F_5,90_=1.017, p=0.412), no significant main effects (n=10, 7, Veh, THC, respectively). **D-K**: Legends

### Enhanced CS-evoked GCaMP8f signal in the LHb during Δ^9^-THC withdrawal

Peak GCaMP8f signals were also measured during 10 pairings of the CS tone with footshock during FC1 and FC2 in the Veh and Δ^9^-THC withdrawal groups (**Fig. 1B-E**). GCaMP8f responses during the first 1 sec of CS were small or absent on the first trials but progressively increased with additional CS-US pairings in both treatment groups (**Fig. 1B-E**, FC1, trial main effect, F_9,170_ = 5.388, p<0.0001). Importantly, when collapsed across all FC1 trials both groups showed significant increases in GCaMP8f signal upon CS presentation (One sample t-test versus no change, Veh = t_9_ = 7.21, p<0.0001, Δ^9^-THC = t_9_ = 7.45, p<0.0001, **S1A**), indicating that the CS-US association was encoded by LHb neurons in both groups. Further analysis showed that the CS GCaMP8f signal was significantly larger in the Δ^9^-THC withdrawal group (**Fig. S1B**, collapsed across all trials, unpaired t-test, t_18_ = 2.21, p = 0.046; **Fig. 1D**, 2-way ANOVA, trial x treatment interaction, F_9, 170_ = 7.64, p < 0.0001), and post hoc analysis revealed differences on trials 4, 6, and 10 (Šídák’s, p=.0056, p<.0001, p=.031, respectively). Differences between groups were less apparent during FC2 (**Fig. 1E**) with no significant trial x treatment interaction (F_1, 140_ = 0.854, p = 0.568), but a significant treatment effect was observed (F_1, 140_ = 4.19, p = 0.043). The results suggest that LHb neurons encode the association between an aversive US and a neutral CS, and that this association is stronger during Δ^9^-THC withdrawal.

### Extinction of LHb CS-evoked GCaMP8f signals during Δ^9^-THC withdrawal

One day after FC2 extinction of the CS-US association was assessed across 2 daily sessions of 20 CS presentations (Ext1, Ext2), with 3 trials averaged into 6 trial blocks (**Fig. 1A, 1F**). The CS-associated calcium signals decreased across Ext1-2 in both withdrawal groups (Ext 1, trial main effect, F_5,102_ = 6.285, p<0.0001, **Fig. 1F-G**), and there was a significant main effect of treatment (F_1,102_ = 5.844, p=0.0174), but no trial x treatment interaction during Ext1 (F_5,102_ = 0.995, p = 0.425). During Ext2 a significant reduction in GCaMP8f signal across trials was observed (trial main effect, F_5,102_ = 5.113, p=0.0003), and the calcium signal was significantly smaller in the Δ^9^-THC withdrawal group on later extinction trials (**Fig. 1G**, treatment main effect, F_1,102_ = 3.95, p=0.0496; unpaired t-test, collapsed across all trials, t_38_ = 2.247, p = 0.0305). However, the trial x treatment interaction was not significant for Ext2 (F_5,102_ = 0.2044, p=0.960). Thus, extinction of the CS-US association progressed over 2 sessions and there was a greater response in the CS-evoked calcium signal during Ext1 in the Δ^9^-THC withdrawal group.

Seven days following Ext2, a CS probe test was conducted in a novel environment to measure generalization to a separate context (**Fig. 1A**, **1H**). The GCaMP8s signal from Δ^9^-THC withdrawal animals (12 days after drug cessation), were significantly larger than the Veh group (trial x treatment interaction, F_5,90_ = 4.80, p=0.0006), with differences observed at the pooled trial block 1-3 (**Fig. 1H**, trials 1-3 pooled, Šídák’s post-hoc, p = 0.0007).

### Increased freezing response to CS during Δ^9^-THC withdrawal during extinction

Behavioral assessment of FC recall was conducted by presenting the CS in the absence of the US following FC1-2 (**Fig. 1I-K**). Although there was no significant interaction between withdrawal condition (Δ^9^-THC or Veh) and trial (F_5,132_=1.52, p=0.188) during Ext1, there were significant main effects of treatment (F_5,132_=11.92, p=0.0007) and trial (F_5,132_=2.82, p=0.019), with post hoc significance on trial blocks 1-3, and 4-6 (Sidak’s; p = 0.0012, p = 0.0144, respectively), indicating more freezing during early CS presentations in Δ^9^-THC withdrawal rats, compared to Veh controls (**Fig. 1I**). In contrast, during Ext2 there was no significant interaction between withdrawal conditions and trial (F_5,132_=1.35, p=0.246), nor a significant main effect of treatment (F_5,132_=0.667, p=0.416; **Fig. 1I**). However, there was a significant trial main effect (F_5,132_=4.04, p=0.0019), with post hoc significance detected at trial block 1-3 (p=0.010, Sidak’s; **Fig. 1I**), suggesting increased freezing in the Δ^9^-THC withdrawal group during the first trial block. To assess fear generalization to another context, a probe test of CS presentations was performed 1 week following Ext2 in a “novel environment” in which the floor of the operant chamber was covered with smooth opaque Plexiglas. Although no significant trial x treatment interaction (F_5,90_=1.017, p=0.412), nor significant main effects were observed, there was a trend toward increased freezing in the Δ^9^-THC withdrawal group during trial blocks 1-3, and 4-6 (**Fig. 1K**).

### LHb excitability is increased, and Inhibitory synaptic transmission impaired during withdrawal from Δ^9^-THC

To determine a potential mechanism for the increased response of LHb neurons to aversive stimuli during Δ^9^-THC withdrawal we used *in vitro* whole-cell electrophysiology in brain slices. Three groups of rats were used; each received 14 daily i.p. injections of either Veh, Δ^9^-THC (5 mg/kg) alone, or the CB1R antagonist AM251 (1 mg/kg), followed 30 min later by injection of 5 mg/kg Δ^9^-THC (Veh, THC, THC+AM groups). Twenty-four hours after the final injection, the synaptic and cellular properties of LHb neurons were assessed.

### Impaired synaptic inhibition of LHb during Δ^9^-THC withdrawal

To determine effects of Δ^9^-THC withdrawal on the integration of excitatory and inhibitory inputs to LHb neurons, we measured glutamatergic EPSCs and GABAergic IPSCs in cells from each of the three treatment groups. Electrically-evoked IPSCs (eIPSCs) were recorded at a holding potential (Vh) of 0 mV, and electrically-evoked EPSCs (eEPSCs) at Vh = -70 mV, in the same LHb cells (**Fig. 2B**). Input-output (I-O) curves plotting the relationship between stimulus intensity and synaptic response amplitude showed that GABAergic eIPSCs were significantly reduced across a range of stimulus intensities in the Δ^9^-THC withdrawal group, compared to the Veh and AM251+THC groups (**Fig. 2C**, 2-way RM-ANOVA, stimulus intensity x treatment interaction, F_36,612_ = 4.255, p < 0.0001, treatment main effect, F_2,34_ = 8.462, p = 0.0010). In contrast, glutamatergic eEPSCs did not differ among the 3 treatment conditions (**Fig. 2D**, 2-way RM-ANOVA, stimulus x treatment interaction = F_36,612_ = 0.737, p = 0.871). To determine whether the deficit in LHb inhibitory transmission during Δ^9^-THC withdrawal resulted from diminished GABA release probability, spontaneously occurring IPSCs (sIPSCs) were measured in LHb neurons from each of the treatment groups. The frequency of sIPSCs was significantly lower in LHb neurons from Δ^9^-THC withdrawal rats, and this was prevented by in vivo AM251 co-administration with the phytocannabinoid (**Fig. 2F-G**, 1-way ANOVA, F_2,46_ = 6.254, p = 0.0040, Tukey’s post-hoc test, Δ^9^-THC vs. Veh, p = 0.0101, Veh vs. THC+AM, p = 0.983). In contrast, sIPSC amplitudes did not differ among the 3 treatment conditions (**Fig. 2H**, 1-way ANOVA, F_2, 46_ = 0.28, p = 0.758). Spontaneous EPSCs from LHb neurons obtained from rats in each of the 3 treatment groups were also measured, and neither sEPSC frequency nor amplitude differed among these groups (**Fig. 2I-K**, 1-way ANOVA, sEPSC frequency, F_2,30_ = 0.3293, p =0.722; sEPSCs amplitude, F_2,30_ = 1.13, p = 0.336, n = 11 neurons in each group).

**Figure 2.**
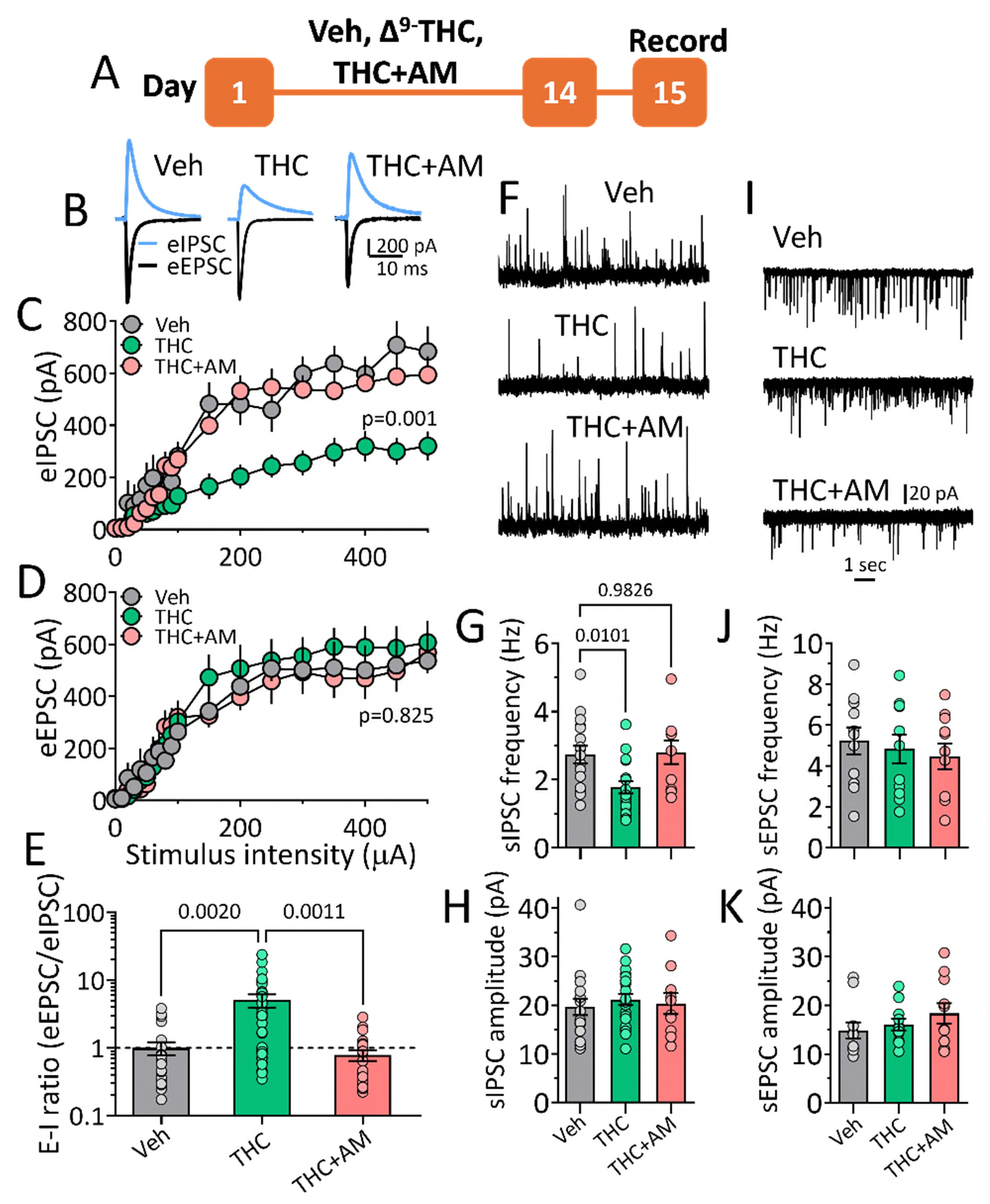
Disruption of E-I balance of synaptic input to LHb neurons during Δ^9^-THC withdrawal. A. Experimental timeline. **B.** Averaged electrically-evoked eIPSCs (blue, Vh = 0 mV) and eEPSCs (black, Vh = –70 mV) recorded simultaneously from LHb neurons in rats withdrawn from Veh, THC, or THC+AM treatments. **C.** eIPSC I-O curves show reduced inhibition during THC withdrawal, prevented by AM251 co-treatment. **D.** No treatment effect on eEPSC I-O curves. **E.** THC withdrawal increased the E-I ratio (eEPSC/eIPSC), blocked by AM251 (log y-axis). **F.** Representative sIPSC sweeps. **G.** THC withdrawal decreased sIPSC frequency (p = .010 vs. Veh, Dunnett’s post-hoc), reversed by AM251. **H.** No change in sIPSC amplitude. **I.** Representative sEPSC sweeps. **J–K.** No differences in sEPSC frequency or amplitude. number rats/number neurons: C–D: 4/9 (Veh), 5/15 (THC), 5/13 (THC+AM); E: 7/23, 8/34, 7/23; G–H: 6/17, 6/22, 5/10; J–K: all groups 4/11.

Another useful measure of altered synaptic integration that has translational value due to its use in establishing baseline levels of excitability and their contribution to changes in circuit bias [41] is known as the excitation/inhibition ratio (E-I). Therefore, we calculated E-I ratios from eIPSCs and eEPSCs recorded in the same neurons and observed roughly similar levels of LHb excitation and inhibition in the Veh control group (E-I ratio = 1.0 ± 0.2, n = 23 cells, **Fig 2E**). However, in neurons from in Δ^9^-THC withdrawal rats, the E-I ratio showed a significant 5-fold increase (E-I ratio = 5.05 ± 1.12, n = 33 cells; 1-way ANOVA, F_2,76_ = 9.38, p = 0.0002, **Fig. 2E**), and this was prevented by AM251 pre-exposure in vivo (E-I ratio = 0.787, ± 1.14, n = 23 cells, p = 0.984, Tukey’s post hoc, **Fig. 2E**). Therefore, collectively these data show that that impaired GABAergic inhibition of the LHb during Δ^9^-THC withdrawal was associated with a large increase in excitability.

### Basal forebrain inhibition of the LHb is selectively impaired during Δ^9^-THC withdrawal

Inhibitory control of LHb neurons is largely derived from extrinsic afferents [42], and we recently identified a GABAergic projection originating in the rostral ventral pallidum (VP) and caudal nucleus accumbens shell (NAcs) of the basal forebrain (BF) that provides strong inhibition of LHb neurons and is controlled by CB1Rs [26]. Previous studies also show that distinct afferents to LHb neurons are not uniformly sensitive to CB1R activation [26,36], and that inhibition of LHb neurons derived from ventral tegmental area (VTA) afferents is not altered by cannabinoids [26]. Therefore, to determine whether the diminished GABAergic control of LHb neurons that was observed during Δ^9^-THC withdrawal requires CB1R expression, we compared selectively activated BF and VTA inputs to the LHb during Δ^9^-THC withdrawal. Separate groups of rats received injections of a channelrhodopsin-2 expressing virus (ChR2, AAV5-hSyn-ChR2-eYFP) into either the BF or the VTA and approximately 6 weeks later underwent long-term treatment with Veh, Δ^9^-THC, or AM251+ Δ^9^-THC. Beginning at 24 hours of withdrawal from treatment, photostimulated IPSCs (oIPSCs) were measured in LHb neuron in vitro. Input-output relationships for oIPSCs evoked from BF inputs showed that these synaptic currents were significantly smaller in LHb neurons in the Δ^9^-THC withdrawal group, compared to the Veh, or THC+AM groups (**Fig. 3B**, 2-way ANOVA, treatment x laser (mW) interaction, F_10,160_ = 10.59, p < 0.0001). In contrast, Inhibitory inputs to LHb from VTA were unaltered in the Δ^9^-THC and THC+AM withdrawal groups, compared to the Veh controls (**Fig. 3C**, 2-way ANOVA, treatment x laser power interaction, F_10,135_ = 0.643, p = 0.775). To determine whether the altered I-O relationship at BF to LHb inputs occurred via a presynaptic mechanism, paired oIPSC responses, evoked by 2 rapid photostimulations of ChR2 at several interstimulus intervals (ISI) at both the BF and VTA projections were measured. The ratio of the first and second oIPSC amplitudes was then calculated (paired-pulse ratio, PPR = oIPSC2/oIPSC1, **Fig. 3D**). Mean PPRs evoked by BF afferent stimulation were less than unity in Veh withdrawal neurons, and this measure was significantly larger than Veh in the Δ^9^-THC withdrawal group (**Fig. 3D**, 2-way RM-ANOVA, ISI x treatment interaction, F_8,152_ = 5.638, p < 0.0001), indicating a decrease in release probability.

**Figure 3.**
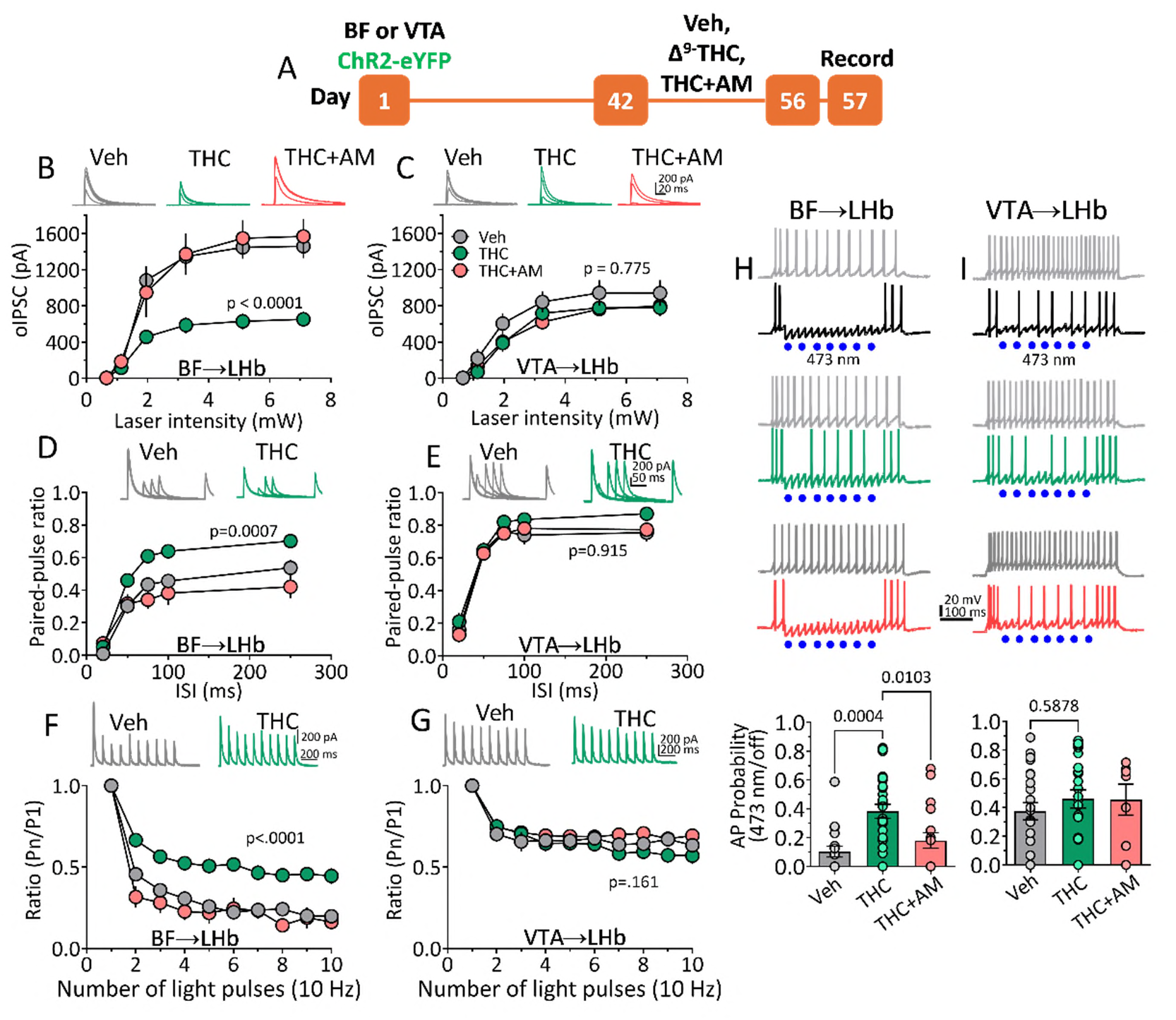
Δ⁹-THC withdrawal alters BF– but not VTA–driven synaptic inhibition of LHb neurons. **A.** Experimental timeline. **B–C.** I-O curves and averaged oIPSCs from BF (**B**) and VTA (**C**) inputs show reduced BF oIPSCs during Δ⁹-THC withdrawal, reversed by AM251; no change at VTA inputs. **D–E.** Paired-pulse ratios (PPRs) across interstimulus intervals. PPR was increased at BF inputs after THC withdrawal (p < 0.0001), indicating decreased release probability; unchanged at VTA inputs. AM251 co-treatment normalized BF PPR. **F–G.** 10 Hz photostimulation revealed reduced short-term synaptic depression at BF inputs during THC withdrawal (**F**), but not at VTA inputs (**G**). BF inputs showed greater depression than VTA under control conditions (p = 0.036), suggesting higher release probability. **H–I.** Current clamp recordings of action potentials (APs) in LHb neurons during current injection with/without photostimulation of BF (**H**) or VTA (**I**) inputs. THC withdrawal reduced BF-mediated AP suppression (p = 0.0003), restored by AM251; no effect on VTA-driven AP suppression. number rats/number neurons: B: 5/12 (Veh), 5/15 (THC), 4/8 (THC+AM); C: 4/9, 5/12, 5/9; D: 6/17, 6/17, 4/7; E: 6/12, 6/12, 5/11; F: 6/11, 7/15, 4/7; G: 6/13, 6/15, 4/7; H: 6/18, 8/27, 6/19; I: 7/24, 7/23, 4/7.

Moreover, AM251 injection preceding Δ^9^-THC prevented the change in PPRs (**Fig. 3D**). In contrast to the changes observed at BF inputs to LHb, PPRs were unaltered at GABAergic VTA inputs to LHb (**Fig. 3E**, 2-way RM-ANOVA, ISI x treatment interaction, F_8,116_ = 0.407, p = 0.915). Differences in release probability are also observed at central synapses during sustained activation of afferents, and this is thought to be relevant to information processing during neuronal oscillations within alpha and theta frequency bands [43]. Therefore, we examined synaptic properties of VTA and BF inputs to LHb neurons during 10 Hz (100 ms ISI) photostimulation and expressed the probability of GABA release as a ratio of oIPSC amplitudes evoked by light pulses 2-10 to those evoked by the first pulse (oIPSC_n_/oIPSC_1_, or P_n_/P_1_ ratio, **Fig. 3F-G**). A comparison between BF- and VTA-LHb inputs showed that baseline probability of GABA release was higher at BF synapses (i.e. more depression of oIPSCs, compared to Veh withdrawal; **Fig. 3F and G**, two tailed t-test, t_18_ = 2.276, p = 0.036). Moreover, an increase in Pn/P1 ratio was observed at BF-LHb synapses from Δ^9^-THC withdrawal rats (**Fig. 3F**, 2-way ANOVA, treatment x pulse number interaction, F_18,288_ = 5.70, p<0.0001), compared to no change in this measure at VTA-LHb synapses (**Fig. 3G**, 2-way ANOVA, treatment x pulse number interaction, F_18,234_ = 1.35, p=0.161). This indicates that GABA release probability is reduced during Δ^9^-THC withdrawal at BF, and not VTA inputs to LHb.

To determine the consequence of reduced inhibitory control during Δ^9^-THC withdrawal on LHb neuron excitability, we examined the effect of photostimulation of BF or VTA afferent input to LHb on the probability of action potential (AP) discharge in current clamp experiments (**Fig. 3H-I**). APs were evoked by 1 sec, 200 pA, square pulses from the patch clamp electrode, either in the absence or presence of 473nm photostimulation of BF or VTA afferents in each of the treatment groups (**Fig. 3H-I**). Photostimulation of either BF or VTA LHb afferents significantly reduced the probability of AP discharge elicited by the depolarization in cells from Veh withdrawal animals (**Fig. 3H-I**, one sample t-test, BF input, t_17_ = 24.16, p < 0.0001; VTA input, t_23_ = 10.39, p < 0.0001). However, the suppression of AP discharge probability by afferent activation was significantly reduced only at BF inputs to LHb neurons in the Δ^9^-THC withdrawal subjects, and this was prevented by AM251 pretreatment in vivo (**Fig. 3H-I**, BF afferents, 1-way ANOVA, F_2,61_ = 9.485, p=0.0003, Veh vs. Δ^9^-THC withdrawal, p = 0.0004, THC vs. THC+AM, p = 0.0103, Tukey’s post-hoc; VTA afferents 1-way ANOVA, F_2,51_ = 0.5390, p=0.587). These results indicate that that Δ^9^-THC withdrawal is associated with a specific impairment of GABA release from BF inputs to the LHb, that this requires CB1Rs, and that inhibitory VTA input to the LHb is unaltered.

### Time course of the impairment of BF inhibition of the LHb during Δ^9^-THC withdrawal

To determine the duration of the impairment of BF inputs to LHb, we compared oIPSC I-O relationships in LHb neurons at 1, 3, 7, 14, or 30 days of Δ^9^-THC withdrawal (**Fig. 4B**). The I-O curves measured at these withdrawal points were significantly different from one another (**Fig. 4B**, 2-way RM ANOVA, light intensity vs. withdrawal day interaction F_25,390_ = 5.509, p<0.0001; withdrawal day main effect, F_5,78_ = 7.322, p<0.0001). Post-hoc (Dunnett’s) analysis indicated I-O curves significantly differed from Veh withdrawal group at 1- and 3-days of Δ^9^-THC withdrawal (p = 0.01, and p = 0.039, respectively).

**Figure 4.**
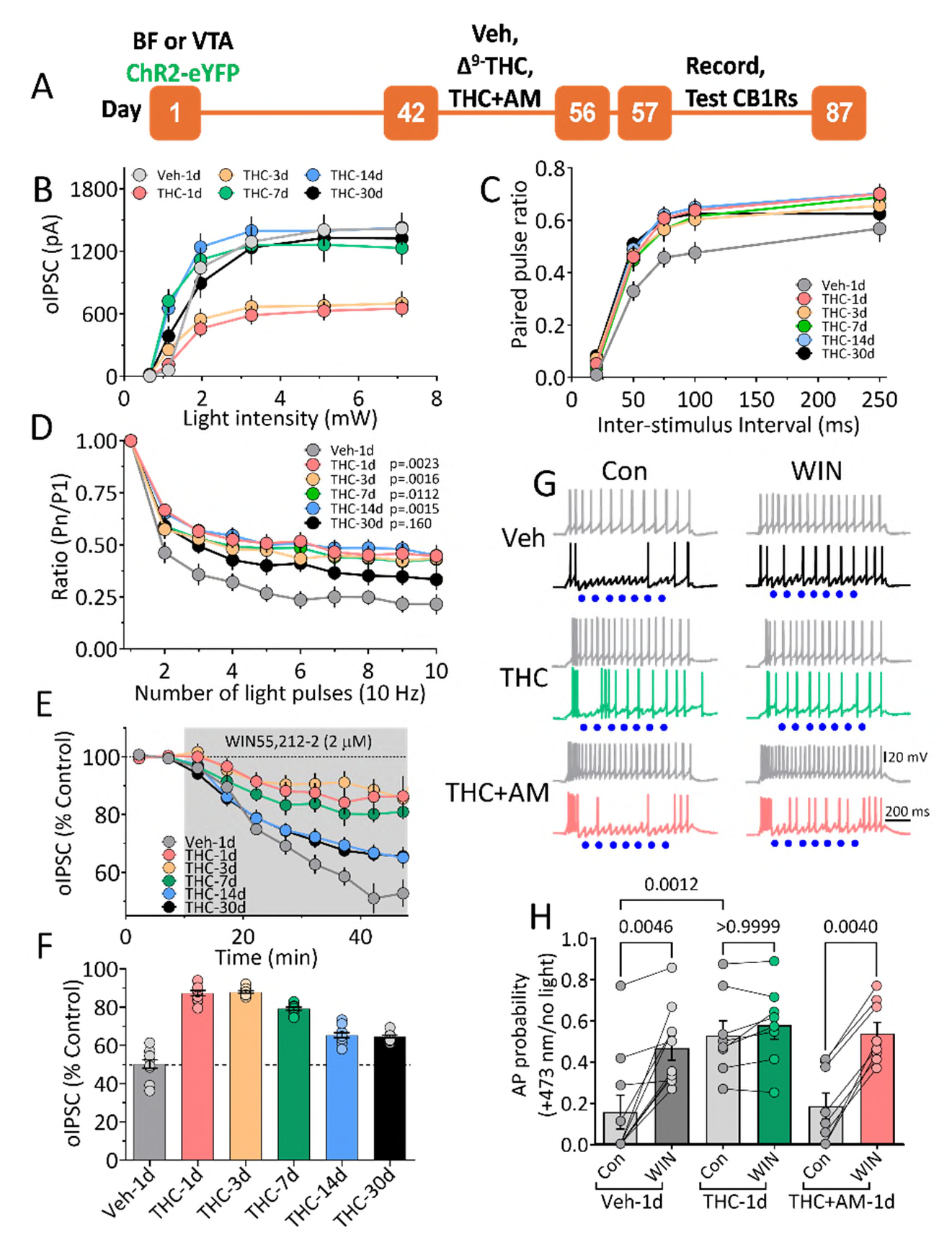
Time course of impaired BF–LHb GABAergic transmission and CB1R desensitization during Δ⁹-THC withdrawal. **A.** Experimental timeline. **B.** I-O curves of photostimulated BF oIPSCs at different days of Δ^9^-THC or Veh withdrawal show reduced inhibition at 1- and 3-days post-withdrawal (vs. Veh-1d; p = 0.01, 0.039, Dunnett post hoc). **C.** PPRs significantly increased at 50, 75, and 100 ms ISIs, at 30d of Δ^9^-THC withdrawal, compared Veh-1d shows enduring synaptic impairment. **D.** 10 Hz photostimulation-induced synaptic depression was reduced at all withdrawal time points except 30d. oIPSCs evoked by pulses 2-10 are expressed as a proportion of oIPSC evoked by the first stimulus. **E–F.** CB1R agonist (WIN, 2 µM) inhibition of BF oIPSCs was diminished throughout 1–30d of Δ⁹-THC withdrawal (p < 0.0001). **G.** Current clamp recordings show reduced BF-mediated inhibition of LHb APs at 1d Δ^9^-THC withdrawal, and loss of WIN effect; AM251 co-treatment restored both. Waveforms show APs evoked by 200 pA, depolarizing current pulses in the absence of BF axon photostimulation (gray waveforms), and during photostimulation (blue dots), in neurons from the Veh-1d (black waveforms), Δ^9^-THC-1d (green waveforms) and THC+AM-1d (red waveforms) groups, before (Control, Con) and during WIN (2µM) application. **H.** Summary of AP probability showing a significant basal increase in the Δ^9^-THC-Con withdrawal group, compared to Veh-Con group (p = 0.0012, Bonferroni post-hoc), and the loss of WIN-induced increase in AP probability in the Δ^9^-THC withdrawal neurons. Note that the change in AP probability was prevented by in vivo treatment with AM251. p values from Bonferroni post-hoc comparisons following a significant (1-way ANOVA (F_5,46_) = 7.682, p<0.0001). number rats/number neurons: B–D: Veh-1d: 5/10–15, THC-1d: 4/14–17, THC-3d: 4/14–15, THC-7d: 4/15, THC-14d: 4/15, THC-30d: 4/14–15; E–F: Veh-1d: 4/9, THC-1d: 4/12, THC-3d: 4/10, THC-7d: 4/11, THC-14d: 4/12, THC-30d: 4/11; H: Veh: 6/18, THC: 8/27, THC+AM: 6/19.

In contrast to the I-O curves of oIPSCs evoked by single light pulses (**Fig. 4B**), the significant increase in PPR seen in the Δ^9^-THC withdrawal group (**Fig. 3D**) did not completely reverse at up to 30 days of Δ^9^-THC withdrawal (withdrawal day main effect, F_5,85_ = 2.96, p = 0.016), with significance observed at the 50, 75 and 100 ms ISIs (**Fig. 4C**, Dunnett’s post-hoc, 30d Δ^9^-THC withdrawal compared to Veh, p = 0.0023, p = 0.018, p = 0.016, respectively). Moreover, the deficit in GABA release was more apparent during sustained activation of BF inputs at 10 Hz (**Fig. 4D**). A significant interaction between the number of stimuli and the number of days of Δ^9^-THC withdrawal was observed (2-way RM-ANOVA, F_45, 711_ = 1.914, p=0.0004), as well as a significant main effect of withdrawal day (F_5,79_ = 3.745, p=0.0043). Moreover, this reduction in the probability of GABA release during 10 Hz photostimulation was significantly smaller, compared to Veh withdrawal neurons, at 1d (THC-1d, Dunnett’s post-hoc, p = 0.0023), 3d (THC-3d, p = 0.00158), 7d (THC-7d, p = 0.0112), and 14d (THC-14dp = 0.0015) of Δ^9^-THC withdrawal, but was not significant at the 30d withdrawal point (THC-30d, p = 0.160; **Fig. 4D**). These results suggest GABA release probability is diminished at BF inputs to the LHb up to 30 days after Δ^9^-THC withdrawal, and this is more evident during higher frequency synaptic activation.

### Time course of CB1R desensitization at BF input to LHb during Δ^9^-THC withdrawal

Our data indicate that GABA release is impaired at BF inputs to LHb during Δ^9^-THC withdrawal, and that this is prevented by co-administration of Δ^9^-THC and AM251 in vivo, indicating that CB1Rs are necessary for the impairment (**Fig. 3**). Therefore, we determined whether CB1R function was altered during Δ^9^-THC withdrawal and measured the duration of these changes using a synthetic cannabinoid agonist.

After collecting baseline oIPSCs evoked by photostimulation of BF afferents to LHb, the agonist WIN 55,212-2 (WIN, 2 µM) was applied to brain slices obtained from rats at the Δ^9^-THC withdrawal time points described above, and the degree of oIPSC inhibition was compared to that observed in the Veh-1d withdrawal group (**Fig. 4E-F**). Consistent with results in drug-naïve rats [26], the mean inhibition of BF afferent-elicited oIPSCs by WIN was 41 ± 3.2% in the Veh-1d withdrawal group (**Fig. 4E-F**). However, in comparison to the Veh-1d group, the response to WIN was significantly smaller in neurons from Δ^9^-THC withdrawal rats at all time points (**Fig. 4F**, 1-way ANOVA, F_5, 60_ = 3.361, p < 0.0001; Dunnett’s post-hoc, Δ^9^-THC withdrawal timepoints 1d-30d, P<0.0001. These data indicate that ability of CB1Rs to inhibit GABA release is impaired for at least 30 days following Δ^9^-THC withdrawal.

### Impaired CB1R control of LHb neuron excitability during Δ^9^-THC withdrawal

To determine how CB1R desensitization might alter cannabinoid control of LHb excitability during Δ^9^-THC withdrawal, the effect of WIN application on the inhibition of AP firing probability caused by photostimulation of BF afferents to LHb neurons in current clamp was examined (**Fig. 4G-H**). Similar to data in **Fig. 3F**, BF afferent inhibition of AP firing was measured during depolarization of LHb neurons via current injection, with and without photostimulation of BF axons (**Fig. 4G**). AP probability was then calculated before, and during application of WIN (2µM) in cells from each of the 1-d withdrawal groups. Photoactivation of BF inputs to LHb neurons from Veh-1d withdrawal rats reduced AP probability, whereas AP discharge significantly increased during WIN application (**Fig. 4G-H**, 1-way ANOVA, F_5,46_ = 7.682, p < 0.0001, p = 0.0046, Bonferroni). In contrast, in neurons from the THC-1d withdrawal group, photoactivation of BF inputs reduced AP probability significantly less than that in the Veh-1d group (**Fig. 4H**, p = 0.0012, compared to control AP probability in Veh-1d group, Bonferroni). Additionally, WIN had no effect on AP probability in cells from THC-1d withdrawal rats (p > 0.999, compared to control AP probability in the THC-1d withdrawal group, Bonferroni), and AM251-co-treatment with in vivo Δ^9^-THC injections prevented the loss of cannabinoid control of AP probability (**Fig. 4H**, p = 0.004, control AP probability vs AP probability during WIN application in the THC+AM-1d group, Bonferroni). These results show that presynaptic CB1Rs inhibit GABA release to modulate the output of LHb neurons, and that this inhibitory control is compromised during Δ^9^-THC withdrawal.

### Passive and active LHb neuron membrane properties are unaltered during Δ^9^-THC withdrawal

Potential changes in the intrinsic excitability of LHb neurons during Δ^9^-THC withdrawal was assessed by examining differences in resting membrane potential (RMP), spontaneous action potential (AP) firing pattern, and the relationship between membrane depolarization and AP discharge in current clamp experiments (**Fig. S2**). There was no effect of treatment on the depolarization-AP relationships (**Fig. S2A**, main effect, 2-way ANOVA, F_2, 72_ = 1.096, p = 0.3399). Similarly, mean LHb neuron RMPs, measured immediately after patch rupture, did not differ among treatment groups (**Fig. S2B** (ANOVA, F_2, 140_ = 0.2903, p=0.7485). LHb neurons exhibit several distinct patterns of spontaneous action potential discharge [44], and we examined potential changes in these patterns at 1d of Δ^9^-THC withdrawal (**Fig. 1C**). Although fewer silent cells (no spontaneous APs immediately after whole-cell access), and more bursting cells were observed in LHb after Δ^9^-THC withdrawal, overall, there was no significant effect of treatment on the pattern of spontaneous cell firing among the treatment groups (**Fig. S1C**, Chi-square value = 5.670, p = 0.1272). These data suggest that Δ^9^-THC withdrawal was not associated with changes in baseline passive and active membrane properties of LHb neurons.

## Discussion

Withdrawal from long-term use of many addictive drugs is associated with negative emotional states that include dysphoria, increased anxiety, elevated responding to aversive stimuli, irritability and increased stress responses [1]. Moreover, avoiding these withdrawal symptoms through continued drug use impedes abstinence [1,45]. As the LHb encodes aversive states [9], can strongly inhibit DA neurons when aversive stimuli are present or when expected rewards are not obtained [8], and because CB1Rs are found within LHb circuits [26,37], we hypothesized that the LHb contributes to cannabis withdrawal-associated negative affective states [46]. To evaluate this, we measured calcium responses with GCaMP8f to track LHb neuron excitability during FC in rats experiencing withdrawal from Δ^9^-THC. Our main results are that the GCaMP8f signal recorded from LHb neurons during presentation of a primary aversive stimulus (footshock, US) is significantly increased during Δ^9^-THC withdrawal, and this was associated with significantly larger calcium signal increases during presentation of a CS (tone) that was paired with the US. Therefore, the data suggest that an increase in LHb neuron responsivity to aversive events facilitates their encoding of previously neutral environmental stimuli that are paired with the aversive event. Additionally, the larger LHb calcium signals in Δ^9^-THC withdrawal rats were also associated with a significant increase in freezing behavior assessed during presentation of the CS during extinction. In support of this behavioral observation, a significant increase in freezing to CS presentation during withdrawal from Δ^9^-THC, administered through cannabis vapor exposure, has also recently been reported [47]. Therefore, the increased response of LHb neurons to aversive stimuli, together with a correlative increase in fear-related behavior, support the idea that the LHb may contribute to the negative emotional state that is observed in humans during cannabis withdrawal.

To examine potential mechanisms for increased responsiveness of LHb neurons to aversive stimuli during Δ^9^-THC withdrawal *in vitro* electrophysiological experiments were conducted. Consistent with the increased LHb GCaMP8f signals *in vivo*, the excitation-inhibition (E-I) ratios of LHb neurons were significantly larger during Δ^9^-THC withdrawal, indicating a shift toward enhanced LHb excitability. Moreover, this shift in E-I balance resulted from reduced synaptic inhibition of LHb neurons, rather than increased excitatory input, which was unchanged. As the increased LHb E-I ratio was prevented by *in vivo* exposure to the CB1R antagonist AM251, prior to each Δ^9^-THC injection, the results indicate that this impairment of inhibitory control involved CB1Rs. Although this represents the first demonstration of increased LHb excitability after Δ^9^-THC exposure, a previous study has shown that mild traumatic brain injury decreases GABA release and also shifts LHb E-I balance toward greater excitation [48], which has been associated with motivational deficits, depression, and anxiety [7,48].

Prior studies from our lab and others show that LHb afferents are not uniformly sensitive to CB1R activation. Of those tested, GABAergic inputs from the BF (VP, NAcs, horizontal diagonal band) and lateral preoptic area (LPO) are inhibited, whereas those arising in VTA or entopeduncular nucleus (EPN) are insensitive to cannabinoids [26,36]. Here we compared specific GABAergic inputs to LHb from BF and VTA during Δ^9^-THC withdrawal using optogenetics and show that GABA release probability was greatly reduced at BF-LHb synapses, whereas GABA release from VTA-LHb afferents was unaffected.

The synaptic impairment at BF-LHb GABAergic synapses following Δ^9^-THC withdrawal also significantly reduced control of LHb neuron firing by these afferents. Therefore, collectively, the results suggest that the GABA release deficit occurs only at CB1R-expressing axons and that the control of LHb excitatory output by BF afferents is impaired during Δ^9^-THC withdrawal.

A GABA release impairment has also been observed at EPN inputs to LHb during cocaine withdrawal, resulting in increased LHb neuron excitability, as well as sensitivity to stress-induced reinstatement of cocaine conditioned place preference [49]. Moreover, the GABA release impairment was associated with decreased expression of the vesicular GABA transporter (VGAT), suggesting that this caused reduced synaptic vesicle concentrations of GABA [49]. Although we did not examine VGAT expression, it is possible that this may represent a common mechanism resulting in impaired control of LHb excitability by GABAergic inputs, as well as the changes in affect that are observed during drug withdrawal.

Our timecourse experiments demonstrated that the impairment of BF-LHb synaptic inhibition persisted for up to 30d following Δ^9^-THC withdrawal, and that the synaptic deficit was more apparent at high frequency synaptic responses. Thus, whereas oIPSCs evoked by single light pulses recovered completely 3-7d after Δ^9^-THC withdrawal, oIPSCs evoked by paired- or 10 Hz-photostimulation were elevated at 30d of withdrawal. These higher frequency photostimulations occur within the range of alpha/theta band neuronal oscillations that are important for temporally binding activity across brain regions during encoding of fearful and anxiogenic stimuli [50–52], and recent studies show that the strength of these oscillations increases in mouse and human LHb in the presence of anxiogenic stimuli [43,53]. Therefore, diminished GABA release probability at these frequencies might be expected to contribute to altered processing of aversive stimuli during Δ^9^-THC withdrawal. Thus, our findings show that this inhibitory synaptic control of LHb neurons is dysregulated for an extended period following Δ^9^-THC withdrawal, and that this may contribute to altered processing of emotional stimuli and enhanced negative affect.

Our study also shows that the recovery timecourse of CB1R inhibition of GABA release parallels that observed for recovery of GABA release probability at BF-LHb inputs. This similarity suggest that perhaps that the stability of GABA release depends upon constitutive CB1R activity that is disrupted by downregulation of CB1R function following repeated Δ^9^-THC exposure [54,55]. In support of this, there is now evidence for tonic CB1R activation by endocannabinoids at both BF and LPO GABAergic inputs to LHb [26,36]. Moreover, a recent study shows, through nanoscale CB1R localization in the hippocampus, that an interaction between CB1R function and maintenance of GABA release probability exists and is dependent upon the proximity of CB1R protein to axon terminal active zones, and that the proximity of these sites is disrupted during Δ^9^-THC withdrawal [56]. Thus, tonic endocannabinoid activation of CB1Rs at axon terminals may contribute to maintaining the relationship between molecular components of neurotransmitter exocytosis to promote synaptic homeostasis, and extended Δ^9^-THC exposure may disrupt this arrangement.

Recently, withdrawal from long-term Δ^9^-THC was reported to decrease LHb neuron AP frequency and change this firing pattern from tonic to phasic in urethane anesthetized rats [28]. Additionally, the acute inhibition of LHb neuron activity by i.v. Δ^9^-THC in this preparation was unchanged [28]. Although the exact reasons for the apparent inhibition of LHb neurons in this study, compared to the enhanced excitability of these cells during withdrawal in our study are unknow, it is notable that we measured LHb calcium signals, as a surrogate for excitability, in awake freely moving rats during presentation of fearful stimuli. Thus, changes in spontaneous LHb neuron firing patterns observed under anesthesia may not reflect that which occurs during presentation of aversive stimuli in awake subjects where sensory input to the LHb is intact. Moreover, as the mechanism for inhibition of LHb neuron firing by i.v. Δ^9^-THC was not identified in this study [28], it remains unclear as to why Δ^9^-THC sensitivity was unaltered during withdrawal.

Our present data support the idea that the LHb may mediate some of the aversive effects of Δ^9^-THC withdrawal that include negative affect, enhanced irritability, and heightened responsiveness to anxiety provoking stimuli [32,45,46]. Thus, GCaMP8f calcium signals are increased during presentation of primary aversive stimuli, and this facilitates LHb neuron encoding of a previously neutral stimulus during Δ^9^-THC withdrawal. At the local circuit level, we show that the LHb E-I balance is strongly shifted to greater excitation and that this is a consequence of reduced probability of GABA release from undefined GABAergic afferents, as well as those arising from BF. As increased LHb neuron excitability contributes to depression [27,57–59], and aversion [10], it is likely that changes in LHb excitability during Δ^9^-THC withdrawal will have implications for cannabis withdrawal and CUD in humans. Additionally, as preclinical data indicate that non-selective inhibition of LHb neurons can reverse the deleterious effects of LHb neuron hyperexcitability on behavior [27,48], we propose that therapeutic interventions designed to limit LHb excitability might be useful to treat several neuropsychiatric conditions including those associated with drug withdrawal.

## Methods

### Subjects

Male and female Long-Evans rats (postnatal day 42-70), purchased from Charles River Laboratories (Rockville, MD, USA) were used for behavioral experiments. For electrophysiological studies male Long-Evans rats of the same age were used. Stereotactic injections of viral constructs were performed at postnatal day 52-57, and *in vitro* electrophysiological experiments were conducted 7-8 weeks later. Animals were housed 2–4 per cage in a temperature and humidity-controlled facility, had *ad libitum* access to food and water, and were housed under standard lighting conditions (lights on 0600 hr, off 1800 hr). All procedures were designed using the “Guide for the Care and Use of Laboratory Animals” [60], and approved by the NIDA-IRP Animal Care and Use Committee. The NIDA-IRP animal facility is accredited by Association for Assessment and Accreditation of Laboratory Animal Care (AAALAC) International.

### Surgery

Rats were initially anesthetized with isoflurane (4%, in 1 L/min O_2_), and then placed the stereotactic apparatus (Kopf Instruments), and the isoflurane concentration lowered to 1%-1.5% (0.2 L/min O_2_). Core temperature was maintained at 37°C using a heating pad. After all surgeries, incisions were closed with absorbable sutures, topical antimicrobial cream was applied to surgical areas, body temperature was maintained until recovery from anesthesia. Rats were injected with a nonsteroidal anti-inflammatory (meloxicam, 1 mg/kg, s.c.), and then returned to their home cage. Weight and health status were monitored for 3 days after surgery.

### Viral construct injections

For electrophysiological experiments, a 10-µl Hamilton syringe with 35 gauge needle, connected to a UltraMicroPump/SYS-Micro4 controller (WPI) was used to inject 0.7 µl of AAV5-hSynhChR2(H134R)-eYFP (5.2×10^12^ virus molecules/ml, University of North Carolina Vector Core), at 100 nl/min, bilaterally into the BF (coordinates from bregma, midline, and level skull, in mm: AP: +1.6; ML: ± 0.8; DV: -6.8), or medial VTA (AP: -5.4; ML: ± 0.2; DV: -8.2, 8° angle). For *in vivo* fiber photometry, 0.3 µl of pGP-AAV-syn-jGCaMP8f-WPRE (GCaMP8f; Addgene, #162376) was infused at 150 µl/min into the right hemisphere LHb (AP: -3.60, ML: ± 0.60, DV: -5.2). The injection needle was left in place for 5 min and slowly withdrawn to promote diffusion into tissue.

### Optical fiber implants

Immediately after LHb GCaMP8f virus injection, 200 µm core diameter optical fibers (Optogenix, Arnesano, Leece, Italy or Thor Labs, USA) were stereotaxically placed using the same LHb coordinates, with optical fiber tips lowered to 0.1 mm above the virus injection location (i.e., -5.1 mm), and anchored to the skull using 4-stainless steel screws and dental cement.

### Behavioral procedures

Three weeks after surgery, rats were gently handled for 2-3 days, prior to LHb GCaMP8f calcium signal testing. To confirm LHb calcium signal recordings, rats were placed in an operant box (Med-Associates, Fairfax, VT), equipped with two cue lights, a white noise generator, a 2900 Hz tone generator, a house fan, digital video camera, and a false floor placed over an electrical grid of stainless-steel rods. Two or three brief air puffs were then delivered through a plastic tube positioned with the tip near the rat’s head. Using video monitoring of the rat’s position, air puffs were implemented when the rat neared the plastic tube. Rats showing increased calcium signals in response to air puffs were then randomly assigned to 1 of the 2 treatment groups. Those animals in which acceptable GCaMP signals were not observed were also randomly assigned to treatment groups to be included in the behavioral freezing analysis.

### Fear conditioning, extinction and probe tests

Two days after the final drug or vehicle injections, rats were individually placed in an operant chamber and connected to an optical fiber tether. After 3–5-minutes of acclimation, the first of 10-fear conditioning (FC1) trials was initiated. Each trial consisted of 5 sec of baseline (Pre-CS), followed by 11 sec of CS (tone) presentation, with the last 0.5 sec of tone coinciding with a mild footshock (unconditioned stimulus, US; 0.65 mA) applied through the grid floor. Each trial then continued for an additional 4 sec (20 sec total time), followed by a 60 sec inter-trial interval, prior to beginning the next trial. This procedure was repeated for 10 trials the following day (FC2). One day after FC2, the first of 20 extinction trials consisting of a 5 sec pre-CS period, 11 sec of CS presentation alone, followed by the 4 sec post-CS period, and a 60 sec inter-trial interval was initiated (Ext1). A second extinction session (Ext2) of 20 trials was conducted 1 day following Ext1. Seven days after Ext2, a probe test consisting of 20 trials of CS alone was conducted in an operant chamber in which the metal grid floor was covered by a smooth plexiglass floor that was painted black. Software (Med-PC) was used to control timing of all behavioral events, and a 28V/+5V TTL adaptor (Med Associates) was connected to the photometry system to mark event timing. For all behavioral experiments, digital video and GCaMP8f calcium signals were recorded for each trial, and behavior later scored offline for time spent freezing (the absence of movement) using the ANY-maze software (Stoelting, Wood Dale, Il, USA).

### Fiber photometry procedures

GCaMP8f signals were collected using a commercially available fiber photometry (FP) system (RZ10x, Tucker-Davis Technologies, Alachua, FL, USA). Two light-emitting diodes (LED) were used; one at 465nm to excite GCaMP8f and one at 405nm (isosbestic control). The timing and intensity of the LED outputs were controlled by the RZ10x system. LED light was filtered and passed through several dichroic mirrors (Fluorescence MiniCube, Doric, Québec, Canada), attached to an optical fiber (200 μm, 0.39 NA, Thorlabs). The MiniCube was also used to direct GCaMP8f emission light from optical fibers to the RZ10x photosensors. The timing and intensity of the light, and recordings of fluorescence signals were set using Synapse software (Tucker-Davis Technologies), with LED intensity set to 40 μW for each channel, measured using an on-board power meter. In order to minimize autofluorescence, cables were photobleached each day prior to initiating recordings. The onset of recording was set to begin 5 sec prior to presentation of a tone channeled through a speaker within the operant chamber.

### Fiber photometry analysis

Fiber photometry data were analyzed off-line using the Photometry Modular Analysis Tool (pMAT version 1.2: Bruno, et al. ^61^. Fluorescence from the signals and isosbestic channels were acquired at 1017Hz via the TDT RZ10x processor. For analysis, peri-event fluorescence data were collected beginning 5 sec before CS presentation at time = 0 sec and continuing for 4.5 sec after US presentation at time = 10.5 sec (total acquisition time = 20 sec). The *ΔF/F* data were converted to z-scores that were normalized to the signal mean and standard deviation during the first 5 sec of recording, prior to CS presentation. Peri-event plots for the CS-US trials were then constructed around CS presentation and collected for group analyses using z-score peri-event histograms generated by pMAT. All measurements of time are shown relative to CS presentation. Pilot comparisons between *ΔF/F* and z-score data showed no statistical differences in aggregate measurements. Therefore, aggregate z-score data are presented for all comparisons of changes in GCaMP8f signal.

### Brain slice preparation and electrophysiological recording

Brain slices containing the LHb were prepared, and recordings from LHb neurons conducted, as described previously [26]. Unless otherwise indicated, electrophysiological studies were conducted 1d after withdrawal from long-term Δ^9^-THC or Veh treatment.

### Drug Treatment

For behavioral experiments, rats were randomly assigned to 1 of 2 treatment conditions. One group received intraperitoneal (i.p.) injections of Δ^9^-THC (5 mg/kg, emulsified in vehicle consisting of 20% DMSO, 10% Tween80, 70% Saline) once per day for 14d, whereas the other group received the same number of injections of vehicle (Veh) alone. This treatment with Δ^9^-THC in rodents results in plasma levels that are similar to those reported for smoked cannabis humans [62,63]. For electrophysiological studies, rats were randomly assigned to 1 of 3 treatment groups: 14d Veh alone (Veh), 14d Δ^9^-THC alone (THC), or 14d of Δ^9^-THC, 30 min after i.p. injection of the CB1R antagonist, AM251 (THC+AM).

### Histology

Localization of GCaMP8f-green fluorescent protein expression and optical fiber implants were evaluated in frozen coronal sections of brain containing the LHb and surrounding regions. Rats were anesthetized with Isoflurane, and transcardially perfused with phosphate-buffer (PB) and 4% paraformaldehyde (PFA). After perfusion, the brain was removed and post-fixed in 4%PFA for 2 hr, then transferred to 30% sucrose until fully submerged. The brain was then frozen in 2-methylbutane at -50°C and stored at -80°C until further processing. Coronal sections (50 µm) were cut using a cryostat (Leica CM 3050S, Germany). Sections were then washed 3 x 10 min in PB, followed by incubation in blocking solution containing 0.3 % Triton X-100 (T8787, Sigma, USA), 4 % bovine serum albumin (A9647, Sigma, USA), and Sodium Azide (S2002, Sigma, USA) for 1 hr at room temperature. After blocking sections were incubated overnight at 4°C with a primary GFP antibody (mouse anti-GFP-632381, 1:1000, mouse, Takara Bio, USA), diluted in blocking solution. Next, sections were washed with PB (3 × 10 min) and incubated for 2 hr at room temperature with secondary antibody (Alexa Fluor 488, donkey anti-mouse, 715545151, Jackson Laboratories, USA). Sections were washed again with PB (3×10 minutes), mounted on slides and counterstained with DAPI (DAPI Fluoromount, 1798424, Electron Microscopy Sciences, USA). Images were then acquired and processed using a VS-200 scanner microscope (Olympus, Evident Scientific, USA) and further analyzed by VS-200 desktop software (Version 4.2).

### Data Analysis

In electrophysiology experiments, representative evoked traces of synaptic currents are averages of 10 consecutive responses prepared using Clampfit (version 9, Molecular Devices, San Jose, CA, USA), and representative spontaneous synaptic current traces are single 10 sec sweeps collected with WinEDR (v4.1.7., University of Strathclyde, Glasgow, UK). Data spreadsheets were constructed using both GraphPad Prism (v10.4.2 Dotmatics, Boston, MA, USA) and Microsoft Excel.

## Statistics

Statistical analyses were performed using GraphPad and data plotted as mean ± standard error of the mean (SEM). Statistical significance was determined using repeated measures (RM) mixed model, 1-way, or 2-way ANOVA where appropriate. Multiple-comparison post hoc tests (Šídák’s, Tukey’s, or Dunnett’s) were conducted only after significant ANOVA interaction, or main effect was observed. In all cases, the minimum significance level was set at P<0.05 using a two-tailed criterion. For statistical analyses in the electrophysiological experiments, “n” describes the number of neurons from which recordings were made, although the number of rats (n_rats_/n_neurons_) is reported in the figure captions.

### Reagents

DL-AP5, 6,7-Dinitroquinoxaline-2,3-dione (DXQX), DL-2-Amino-5-phosphonopentanoic acid (DL-AP5), (R)-(+)-WIN 55212, and AM251 were purchased from Tocris Bioscience (Bristol, UK). Δ^9^-THC was obtained from the National Institute on Drug Abuse Drug Supply Program (Rockville, MD). Drugs in electrophysiological experiments were dissolved in artificial cerebrospinal fluid (aCSF) and delivered to brain slices via a gravity perfusion system. Drugs were prepared as stock solutions in H_2_O or DMSO and Tween 80 and diluted to the indicated concentrations.

## Data availability

All data generated or analyzed during this study are included in this article or its supplementary files.

## Acknowledgement

This research was supported by the NIH (NIDA) Intramural Research Program.

## Author Contributions

E-KH designed, performed, and analyzed electrophysiological experiments; AZ, DD and CCM designed, performed and analyzed behavioral and fiber photometry experiments and edited the manuscript. DD and AZ performed histological analyses and confirmed virus and optical fiber placements. AFH and AZ participated in the design of the study, performed video analysis of freezing data, and edited the manuscript; CRL designed and supervised the study, analyzed the data, performed statistical analyses, and wrote and edited the manuscript.

---

**Supplemental Figure 1.**
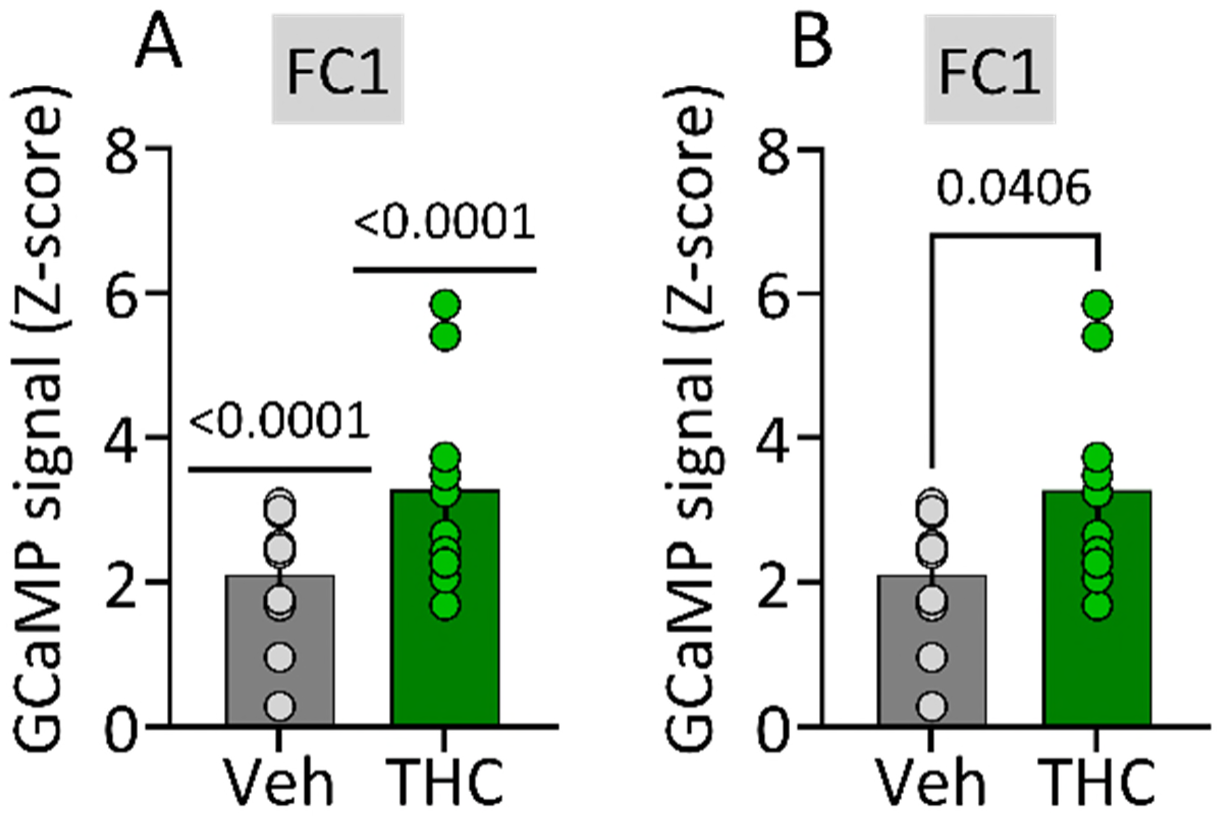
Mean LHb GCaMP8f responses to CS during FC1 pooled across trials 1-10 in Veh and Δ^9^-THC withdrawn rats. All data are from those shown for each trial in **Fig. 1D** (bottom panel). **A**. Results of 1 sample t-test showing a significant increase in GCaMP8f signal z-score to the CS during CS-US pairings. **B.** Results of unpaired t-test showing a significant increase in GCaMP8f signal in the Δ^9^-THC withdrawal group (n’s: Veh = 12, Δ^9^-THC = 7).

**Supplemental Figure 2.**
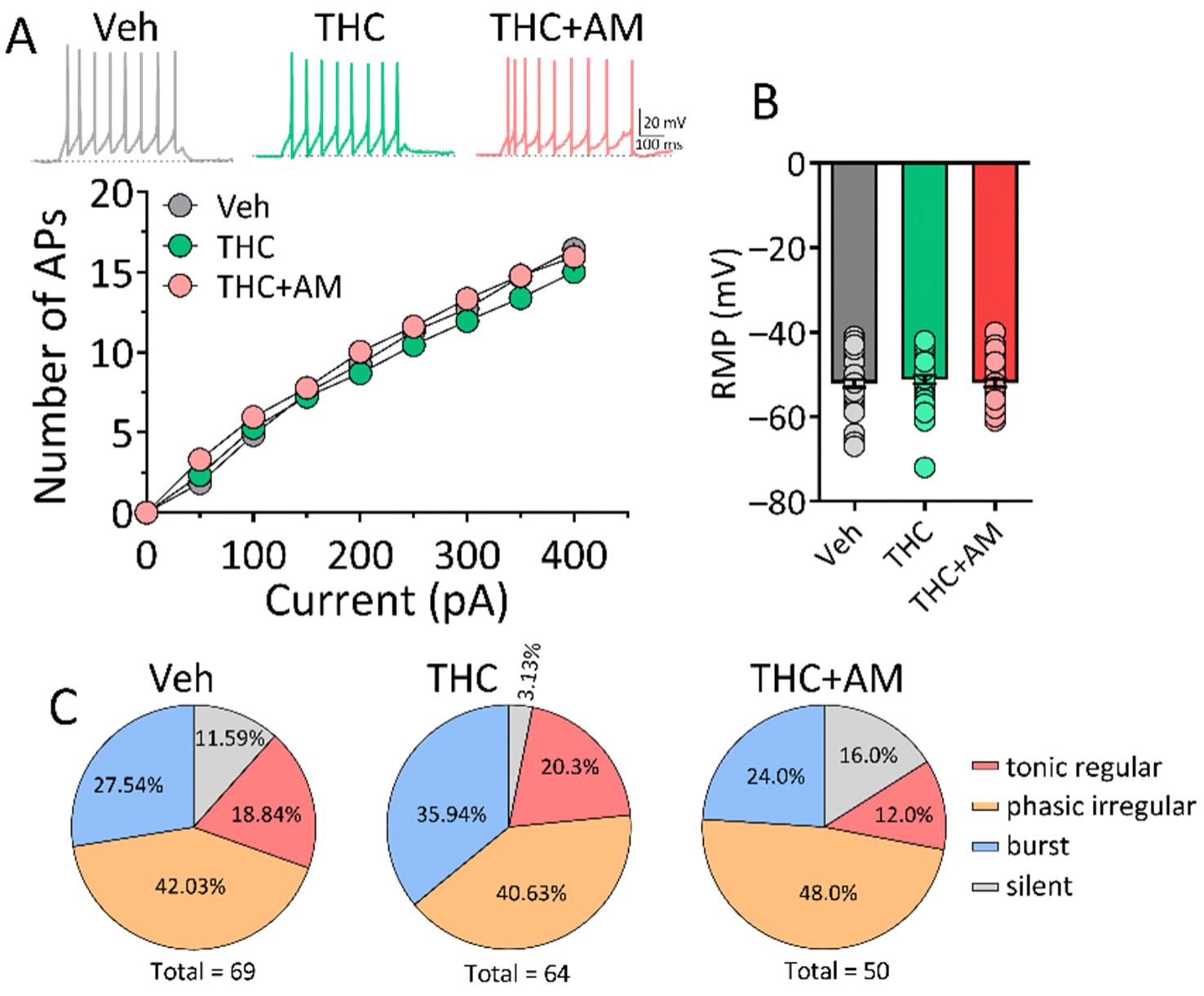
No change in active or passive membrane properties of LHb neurons at 1d Δ^9^-THC withdrawal. **A**. Unaltered relationship between injected current amplitude and AP number in all 1-d withdrawal treatment groups. **B**. No change in resting membrane potential across all withdrawal treatment groups. **C**. No change in spontaneous firing patterns of LHb neurons during withdrawal from all treatment conditions *in vitro* (X^2^ = 5.70, p = 0.1272. n_rats/_n_neurons_: **A-B**, Veh = 10/47, THC = 11/49, THC+AM = 10/47.

